# Network-Based Analysis Identifies Targetable Pathways in Comorbid Type II Diabetes and Neuropsychiatric Disorders

**DOI:** 10.1101/2024.06.25.600630

**Authors:** Anna Onisiforou, Panos Zanos

## Abstract

Comorbid diseases complicate patient outcomes and escalate healthcare costs, necessitating a deeper mechanistic understanding. Neuropsychiatric disorders (NPDs) such as Neurotic Disorder, Major Depression, Bipolar Disorder, Anxiety Disorder, and Schizophrenia significantly exacerbate Type 2 Diabetes Mellitus (DM2), often leading to suboptimal treatment outcomes. The neurobiological underpinnings of this comorbidity remain poorly understood. To address this, we developed a novel pathway-based network computational framework that identifies critical common disease mechanisms between DM2 and the five prevalent NPDs. Our approach involves reconstructing an integrated DM2 ∩ NPDs KEGG pathway network and applying two complementary analytical methods, including the “minimum path to comorbidity” method to identify the shortest pathways fostering comorbid development. This analysis uncovered shared pathways like the PI3K-Akt signaling pathway and highlighted key nodes such as calcium signaling, MAPK, estrogen signaling, and apoptosis pathways. The dysregulation of these pathways likely contributes to the development of DM2-NPDs comorbidity. Our model not only elucidates the intricate molecular interactions driving this comorbidity but also identifies promising therapeutic targets, paving the way for innovative treatment strategies. This framework can be adapted to study other complex comorbid conditions, offering broad implications for improving patient care.

## Introduction

Comorbidity, defined as the presence of two or more diseases in the same individual, is often due to casual association, chance or selection bias (1). The presence of comorbid diseases is associated with worse patient outcome, more complicated treatments and increased healthcare costs (1, 2). Thus, understanding the etiology of comorbid diseases is essential for appropriate treatment of comorbidities and the prevention of their emergence.

Type 2 Diabetes mellitus (DM2) is a chronic metabolic disorder affecting approximately 6.28% of the world’s population, corresponding to 462 million people (3). Projections indicate that the disease will affect 700 million people by 2045 (4). DM2 is characterized by insulin resistance and hyperglycemia, accounting for 90% of all diabetes cases (5). It occurs when pancreatic islet β cells fail to produce sufficient insulin to maintain normal glucose metabolism (6, 7). The origin of DM2 is multifactorial, involving the complex interaction of genetic, environmental and lifestyle factors (8). Although DM2 most often develops in people older than 45, in recent years, there has been a rise in incidence among younger populations due to obesity, increased food intake, and lack of physical activity (9, 10). The risk of DM2 increases with age, with approximately one-third of the individuals suffering from DM2 being above 50 years old (3).

In addition to the peripheral symptoms, DM2 patients frequently experience emotional and behavioral symptoms, such as anxiety and depression, which exacerbate disease severity (11). DM2 is associated with a higher incidence of both major and minor depressive disorders (12). A bidirectional relationship seems to exist between DM2 and depression, where individuals with DM2 have nearly double the risk of developing depression, and individuals with depression have an increased risk of developing DM2 (13). Moreover, individuals with bipolar disorder (BD) (14) and schizophrenia (15) have three to five times higher risk of developing comorbid DM2. Overall, current evidence suggests that individuals with neuropsychiatric disorders (NPDs) have an increased risk of developing DM2, and *vice versa*. Comorbid NPDs in DM2 patients are linked to an impaired quality of life (16), reduced treatment adherence (17), decreased control over glycemia (18), and poorer treatment prognosis (19, 20).

Despite the widespread acceptance of comorbidity between DM2 and NPDs and some evidence of shared pathophysiology, the exact mechanisms underlying this comorbidity are not fully understood. Currently, no available pharmacological interventions are specifically available for treating comorbid DM2 and NPDs. In fact, in some cases, treatments for DM2 can worsen NPDs, and *vice versa* (21–24). Therefore, there is an urgent need to understand the mechanisms facilitating the emergence of comorbid DM2 and NPDs.

Our study addresses this need by applying network-based approaches to a unique combination of disorders—DM2 with MDD, ND, AD, BD, and Schizophrenia. Unlike previous studies that have focused on isolated conditions or different comorbidity pairs, our research provides novel insights into the shared molecular pathways and potential therapeutic targets for this specific set of comorbid conditions.

Network-based approaches, including protein-to-protein interaction (PPI) networks, have been instrumental in unravelling common and shared pathological mechanisms between comorbid diseases (25–27). Barabasi’s work on network medicine has provided a comprehensive framework for understanding the complexity of molecular interactions in human diseases (28). Brunak et al. have contributed to the field with their studies on integrated molecular networks and the identification of disease-promoting genes through a systematic, large-scale analysis of human protein complexes (29). However, no studies have specifically focused on the comorbidity between DM2 and a spectrum of NPDs including Major Depressive Disorder (MDD), Neurotic Disorder (ND), Anxiety Disorder (AD), Bipolar Disorder (BD), and Schizophrenia. Previous research has typically investigated DM2 comorbidities with individual disorders like Alzheimer’s Disease or studied NPDs in isolation using network-based approaches. For example, in the context of comorbid DM2 related-diseases, some studies have analyzed transcriptomic data, using network-based approaches, and identified common pathophysiological mechanisms between DM2 and Alzheimer’s Disease (27, 30). Additionally, network-based methodologies have been utilized to study NPDs, including unravelling pathogenic mechanisms that can lead to their development (31) and identifying existing drugs that can be potentially used for their treatment (32). Pathway-based analysis methods have also significantly advanced our understanding of complex diseases as they provide insights into their molecular mechanisms through the exploration of large-omics data (33).

Here, we developed and applied a pathway-based network computational framework (see **Figure 1**) that enables to unravel key pathological mechanisms that could facilitate the development of comorbidities, specifically DM2 and NPDs. To the best of our knowledge, no prior studies have used network-based approaches specifically to study the emergence of this comorbidity. Our study is novel in its comprehensive approach to understanding the shared pathophysiological mechanisms between DM2 and these five prevalent NPDs concurrently, providing new insights into the intricate molecular interactions that drive these comorbid conditions. First, we identified common disease pathways shared between DM2 and all five NPDs, ND, MDD, AD, Schizophrenia and BD, through the reconstruction and analysis of five DM2-NPDs comorbidity protein-protein interaction (PPI) networks. These particular NPDs were selected because they frequently occur alongside DM2. The shared pathways between DM2 and these NPD were then used to reconstruct the DM2 ∩ NPDs KEGG pathway-to-pathway network. Furthermore, we employed two complementary strategies to analyze the DM2 ∩ NPDs KEGG pathway-pathway network and pinpoint key pathways contributing to this comorbidity. The initial approach focused on identifying the top 10 high centrality pathways that could potentially exert system comorbid effects. Finally, we devised the ‘minimum path to comorbidity’ method, leveraging graph theory techniques to isolate the shortest path that might facilitate the development of comorbid DM2 and NPDs. This approach highlighted key disease pathways that functionally interact and are in close proximity to the reference pathways representing DM2 and NPDs, suggesting that targeting these pathways pharmacologically could have a higher therapeutic impact.

**Figure 1:**
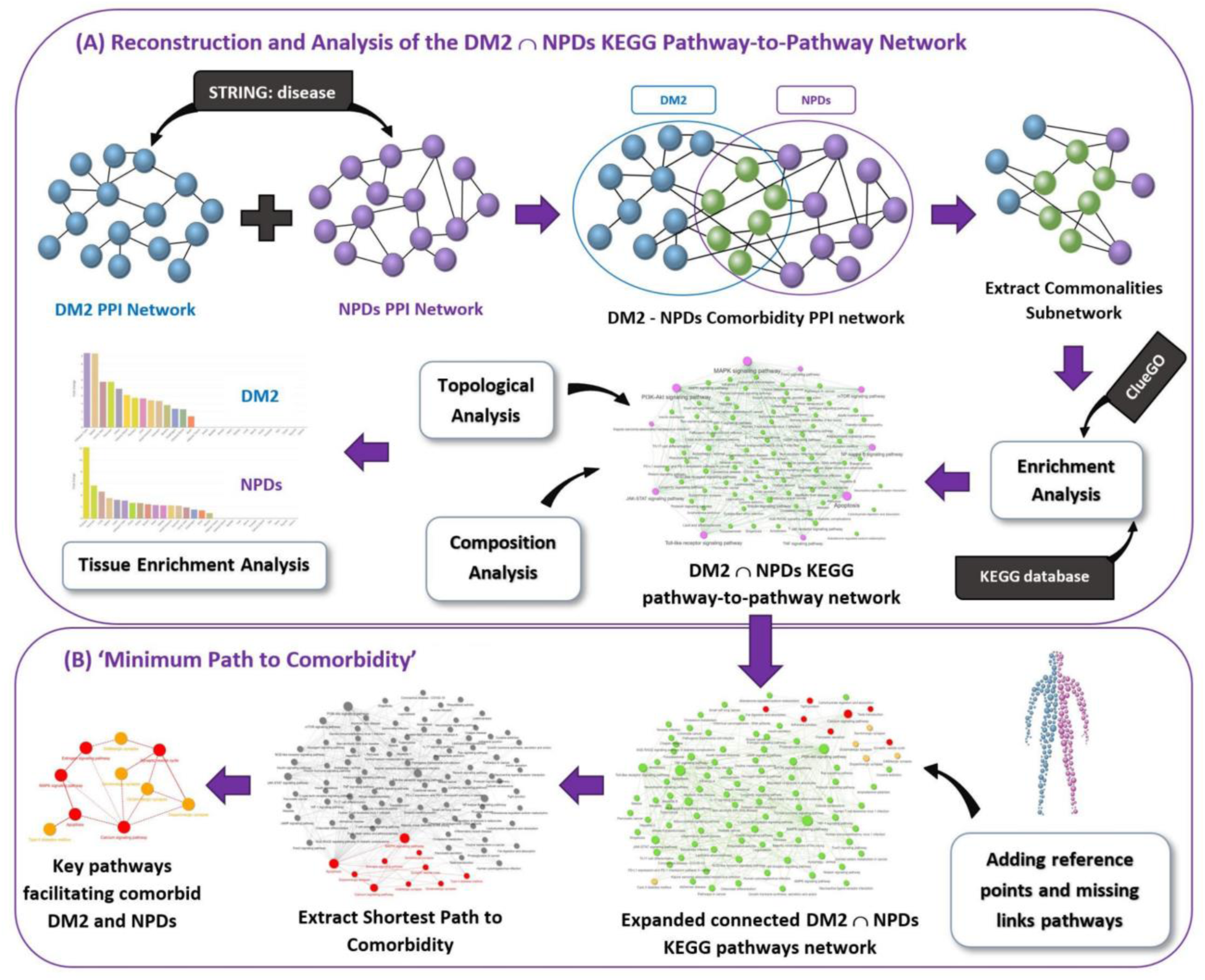
Schematic illustration of the methodology used in this paper. The aim was to isolate key pathways contributing to the development of comorbid DM2 and NPDs. **(A)** We represent the various data sources and the methodology used to reconstruct and analyze the DM2 ∩ NPDs KEGG pathway-pathway network. Topological analysis was employed to identify the top 10 high centrality pathways, and composition analysis was performed to determine the subclass to which the common pathways belong. **(B)** We also devised the ‘minimum path to comorbidity’ method, which allows us to isolate the shortest path that facilitates the development of comorbid DM2 and NPDs.

## Methods

### Reconstruction of the Integrated DM2-NPDs Comorbidity Protein-to-Protein Interaction (PPI) Networks

To identify common pathological mechanisms between DM2 and each of the five NPDs, we reconstructed five DM2-NPDs comorbidity PPI networks (**Table 1)**. Following established studies (31, 34–36), we utilized the *STRING disease* app in Cytoscape to collect the top 200 disease-associated proteins, ranked by the highest disease association score, for each of the six conditions: Type 2 Diabetes mellitus (DM2) (DOID:9352), Major depressive disorder (MDD) (DOID:1470), Neurotic Disorder (ND) (DOID:4964), Anxiety disorder (AD) (DOID:2030), Bipolar disorder (BD) (DOID:3312), and Schizophrenia (DOID:5419). The *STRING disease* app obtains its data from the DISEASES database (37), which collects gene-disease associations from various types of evidence, including automatic text mining, manually curated databases like UniProt Knowledgebase (UniProtKB), genome-wide association studies, and cancer mutation data. These associations are then unified and assigned a confidence score, with 5 stars indicating high confidence and 1 star indicating low confidence in the association being a true positive. Focusing on the top 200 high ranking proteins enables the prioritization of proteins strongly implicated in the diseases of interest, facilitating the identification of biological processes closely associated with disease progression (38).

**Table 1:** Characteristics of the DM2-NPDs comorbidity PPI networks and commonalities subnetworks.

| DM2-NPDs comorbidity PPI networks | Nodes | Edges | Common disease-associated proteins ($DM2 \cap NPDs$) | Commonalities subnetwork human proteins |
| --- | --- | --- | --- | --- |
| DM2 and AD | 368 | 1610 | 32 | 197 |
| DM2 and MDD | 381 | 1558 | 19 | 177 |
| DM2 and Schizophrenia | 379 | 1610 | 21 | 181 |
| DM2 and BD | 381 | 1588 | 19 | 170 |
| DM2 and Neurotic Disorder | 360 | 1531 | 40 | 206 |

To reconstruct the five DM2-NPDs comorbidity PPI networks, we merged the DM2 PPI network with each of the five NPDs PPI networks using the “merged” function in Cytoscape (see **Figure 1A**). The confidence cut-off score for the PPI was set at 0.8. This score is determined based on the nature and quality of the supporting evidence for the PPIs, ranging from 0 (indicating low confidence) to 1.0 (indicating high confidence). Therefore, the higher the score, the greater the likelihood that the PPIs are true positives (39). It’s worth noting that a recommended cut-off for high confidence is above 0.7 (40). As a result, a stronger cut-off of 0.8 was selected.

### Commonalities Subnetwork Extraction and Enrichment Analysis

We, then extracted the commonalities subnetwork from each DM2-NPDs comorbidity PPI network. These subnetworks include common disease-associated proteins between DM2 and each NPD, along with their first neighbors. The number of common disease-associated proteins between DM2 and each NPDs found from each comorbidity network, along with the number of human proteins contained in each extracted commonalities subnetwork, is listed in **Table 1**. Using the isolated human proteins from each subnetwork, we then performed enrichment analysis using the Kyoto Encyclopedia of Genes and Genomes (KEGG) database (41). Pathway enrichment analysis allows us to gain mechanistic insight associated with a list of proteins (42). Enrichment analysis was conducted in the ClueGO app (43) in Cytoscape, utilizing the KEGG database. Only statistically significant enriched terms with an adjusted *p*-value ≤ 0.05 (corrected with Bonferroni step-down) were retained.

### Reconstruction and Analysis of the DM2 ∩ NPDs KEGG Pathway-to-Pathway Network

#### (a) Integrated DM2 ∩ NPDs KEGG Pathway-Pathway Network

To identify common pathological pathways between DM2 and all the NPDs included in our analysis, we reconstructed an integrated DM2 ∩ NPDs KEGG pathway-pathway network. In this network, nodes represent pathways, and edges denote functional relationships between these pathways. To reconstruct the integrated network, we first compared the isolated significantly enriched KEGG pathways found between DM2 and each of the five NPDs using enrichment analysis. This allowed us to identify the KEGG pathways involved in DM2 that are also found in all five NPDs. Overall, we identified 87 KEGG pathways shared between DM2 and all five NPDs.

To ensure that the identified 87 common pathways do not contain any false positive results of pathways containing only DM2- or NPD-associated genes, we checked that all the pathways contained at least one disease-associated gene for each condition. Using this method, we did not identify any false positive results, as all of the 87 common pathways were found to contain at least two genes associated with each condition.

To reconstruct the integrated DM2 ∩ NPDs KEGG pathway-pathway network and create functional interactions between the 87 common pathways, we utilized the KEGGREST package (44) in R to parse the KEGG database (41). We parsed each of the 353 KEGG pathways (Homo Sapiens) entries to extract information of the functional relationships of each pathway with other pathways. We then combined all the interactions obtained, which resulted into 1914 functional pathway-to-pathway interactions. Functional relationships between pathways, represent the interconnectedness and communication that occurs between different pathways to accomplish complex physiological processes, such as where components from one pathway influence the activity or regulation of components in another pathway. We then isolated the functional relationships between the 87 common pathways and reconstruct a direct DM2 ∩ NPDs KEGG pathway-pathway network, composed of 87 nodes and 328 edges. Therefore, these 87 pathways represent shared biological mechanisms that concurrently play a role in both diseases, with the edge interactions denoting functional relationships between the pathways

#### (b) Pinpointing Essential Comorbidity Disease Communicator Nodes

After reconstructing the DM2 ∩ NPDs KEGG pathway-pathway network, we utilized the igraph package (45) in R to conduct network analysis and identify high centrality pathways, specifically focusing on nodes that function as hubs within the network. These hub nodes are crucial for facilitating interactions between pathways and ensuring the flow of information within the network. As a result, they exert systemic comorbid effects by functionally interacting with multiple pathways. Hence, they serve as key comorbidity disease communicator nodes (34) and play a crucial role in promoting the emergence of comorbid DM2 and NPDs. To pinpoint these essential nodes, we isolated the top 10 nodes in the network based on their highest hub scores, which serve as indicators of high centrality.

#### (c) Composition Analysis and Tissue-Specificity Enrichment Analysis of DM2-NPDs Interactions

In addition, we conducted composition analysis on the DM2 ∩ NPDs KEGG pathways network to determine the subclasses to which the 87 common disease pathways belong. To achieve this, we used the KEGGREST package (44) in R to extract the subclass classification of each of the 87 pathways based on the KEGG database. This analysis offers valuable insights into the categorization of pathways and their functional relevance in the context of comorbid DM2 and NPDs.

Moreover, in order to identify potential overlapping tissues where the dysregulation of the 87 identified pathways could occur between DM2 and NPDs, we performed tissue-specific gene enrichment analysis. This analysis involved using the disease-associated genes identified to participate in these pathways from each condition. We performed the analysis using the TissueEnrich web application (46), and chose the GTEx database (47). The GTEx database provides the most comprehensive information on normal tissue expression across 56 distinct tissues. It is important to note that in the TissueEnrich application, samples from sub-tissues are combined into broader categories. For example, different regions of the brain are grouped under the term “brain”, resulting in 29 human tissue categories. To identify tissue-specific genes, we applied the “tissue enriched” criterion, which defines genes as tissue-specific if their expression levels are at least five times higher in a particular tissue compared to all other tissues. In addition, we employed the fold-change test to determine the statistical significance of the tissue-specific genes.

### The ‘Minimum Path to Comorbidity’

To isolate the shortest path that may facilitate the development of comorbid DM2 and NPDs, we developed a method termed the ‘minimum path to comorbidity’ (see **Figure 1B**). This approach leverages graph theory methods, specifically the shortest path method, to isolate the ‘minimum path to comorbidity’, thereby uncovering key routes that contribute to the emergence of comorbid DM2 and NPDs.

#### (a) Highlighting and Adding Missing Reference Points on the DM2 ∩ NPDs KEGG Pathway-Pathway Network

To determine the “minimum path to comorbidity” between DM2 and NPDs, we first selected five KEGG pathways as reference points: (i) the Type II diabetes mellitus (hsa04930) pathway, representing DM2, and (ii) four KEGG pathways—Dopaminergic synapse (hsa04728), Glutamatergic synapse (hsa04724), Serotonergic synapse (hsa04726), and GABAergic synapse (hsa04727)—representing NPDs. While specific disease pathways for the five NPDs under investigation are not available in the KEGG database, we selected these four pathways to collectively represent the NPDs “pathway,” as they play crucial roles in the development of all NPDs in our analysis (48–51). Although other KEGG pathways might also be linked to these NPDs, these selected pathways serve as reference points because they are crucial targets of current pharmacological interventions used for the treatment of these NPDs. For example, selective serotonin reuptake inhibitors, such as fluoxetine and sertraline, that are commonly prescribed for the treatment of MDD, BD, ND and AD, target the serotonin system. Moreover, the antidepressant ketamine used for treatment-resistant depression, exerts its effects by modulating glutamate neurotransmission (52, 53). Additionally, benzodiazepines, such as lorazepam, used to treat MDD, ND and AD, enhance GABAergic activity in the brain (54). Furthermore, atypical antipsychotic drugs, such as risperidone, used for the treatment of BD, MDD and Schizophrenia affect both the dopaminergic and serotonergic systems (55). All the aforementioned highlight the importance of the selected pathways in these NPDs.

Our approach works by first highlighting the selected reference points on the DM2 ∩ NPDs KEGG pathway-pathway network in yellow. However, only two reference points, the Dopaminergic synapse and Type II diabetes mellitus pathways, were initially present on the network. The absence of the other reference points highlights a limitation of enrichment analysis. Although the Glutamatergic, Serotonergic, and GABAergic synapse pathways play crucial roles in the pathogenesis of these NPDs, they were not identified as statistically significant during the enrichment analysis. To overcome this limitation, our methodology adds the three missing reference points - Glutamatergic synapse, Serotonergic synapse, and GABAergic synapse - to the network. To add the missing reference points, the algorithm calculates all the shortest paths between the reference points and all the nodes on the DM2 ∩ NPDs KEGG pathway-pathway network, using the functional relationships of all 353 KEGG pathways collected from KEGG database. Subsequently, it identifies the shortest path with the smallest length for each missing reference point and extracts the relevant edge interactions between the reference points and the pathways on the network. In the case where no direct interactions exist between the missing reference points and the nodes on the network, the algorithm introduces additional missing nodes and edge interactions that are required to connect the missing reference points with the rest of the network. In addition, if multiple shortest paths with the smallest length exist, the algorithm adds the interactions for all of these paths to the network.

#### (b) Adding Missing Pathways on the Expanded DM2 ∩ NPDs KEGG Pathway-Pathway Network

Moreover, the algorithm aims to create a fully connected network, where all nodes are connected with each other, so it identifies all nodes with degree values of 0 and 1 and determines all pairs that exist between these nodes and the remaining nodes on the expanded network. It then calculates all the shortest paths between the pairs using the functional relationships collected from KEGG database. For each node of interest, the algorithm isolates the shortest path with the smallest length. Finally, the algorithm extracts and adds the relevant edges and missing nodes to create a fully connected expanded DM2 ∩ NPDs pathway-pathway network. The reliability of our graph expansion approach in introducing relevant comorbidity disease pathways in a Disease ∩ Disease KEGG pathway-pathway network is further analyzed in ***Supplementary File 1***.

#### (c) Isolating the Minimum Path to DM2 and NPDs Comorbidity

To identify the key pathways that facilitate the development of comorbid DM2 and NPDs, we utilized the connected expanded DM2 ∩ NPDs KEGG pathway-pathway network to isolate the minimum path to DM2 and NPDs comorbidity. This process involved calculating the shortest paths with the smallest length for each of the four pairs that exist between the Type II diabetes mellitus pathway (DM2 reference point) and each of the four pathways (Dopaminergic, Glutamatergic, Serotonergic and GABAergic synapses) that represent the NPDs reference points. We then highlighted the corresponding edges on the fully connected expanded DM2 ∩ NPDs pathway-pathway network. This enabled us to isolate the minimum path between DM2 reference point and the four NPDs reference points.

### Validation of the Importance of the Central and “Shortest Path to Comorbidity” Pathways in DM2 and NPDs Comorbidity Using Prefrontal Cortex Data

To validate the importance of the central and “shortest path to comorbidity” pathways in the context of DM2 and NPDs comorbidity, we conducted an analysis using microarray gene expression data from the prefrontal cortex (PFC). We accessed this data through the Gene Expression Omnibus (GEO) database, a repository of transcriptomic information (56). We sought PFC microarray studies associated with NPDs, DM2, and comorbid DM2 and NPDs. Unfortunately, there were no available datasets for PFC samples from humans with DM2 or datasets from either animal models or humans with comorbid DM2 and NPDs. However, we identified dataset GSE34451, which includes three samples from the PFC of male Goto-Kakizaki rats (a model of Type II diabetes) and three samples from male Wistar rats (control) (57). Additionally, we analyzed dataset GSE12654, which contains human PFC samples from patients with various NPDs (58).

The Limma R package (59), which allows for the identification of differentially expressed genes (DEGs) from microarray experiments, was used to analyze each dataset and identify DEGs between each condition and the control samples. Both datasets were normalized and log_2_ transformed. DEGs with *p*-value <0.05 were considered as statistically significant. Subsequently, pathway enrichment analysis was performed using the ClueGO app (43) in Cytoscape, utilizing the KEGG database, to identify statistically significant enriched pathways with an adjusted *p*-value ≤ 0.05 (corrected with Benjamini-Hochberg) that are related to the DEGs.

## Results

### Comparison between the five NPDs

Comparison between the top 200 disease-associated proteins of the five NPDs (see Methods) using a Venn diagram (60) revealed that they share 43 common disease-associated proteins (**Figure 2**). **Table 2** indicates the major biological processes to which the disease-associated proteins of the NPDs belong.

**Figure 2:**
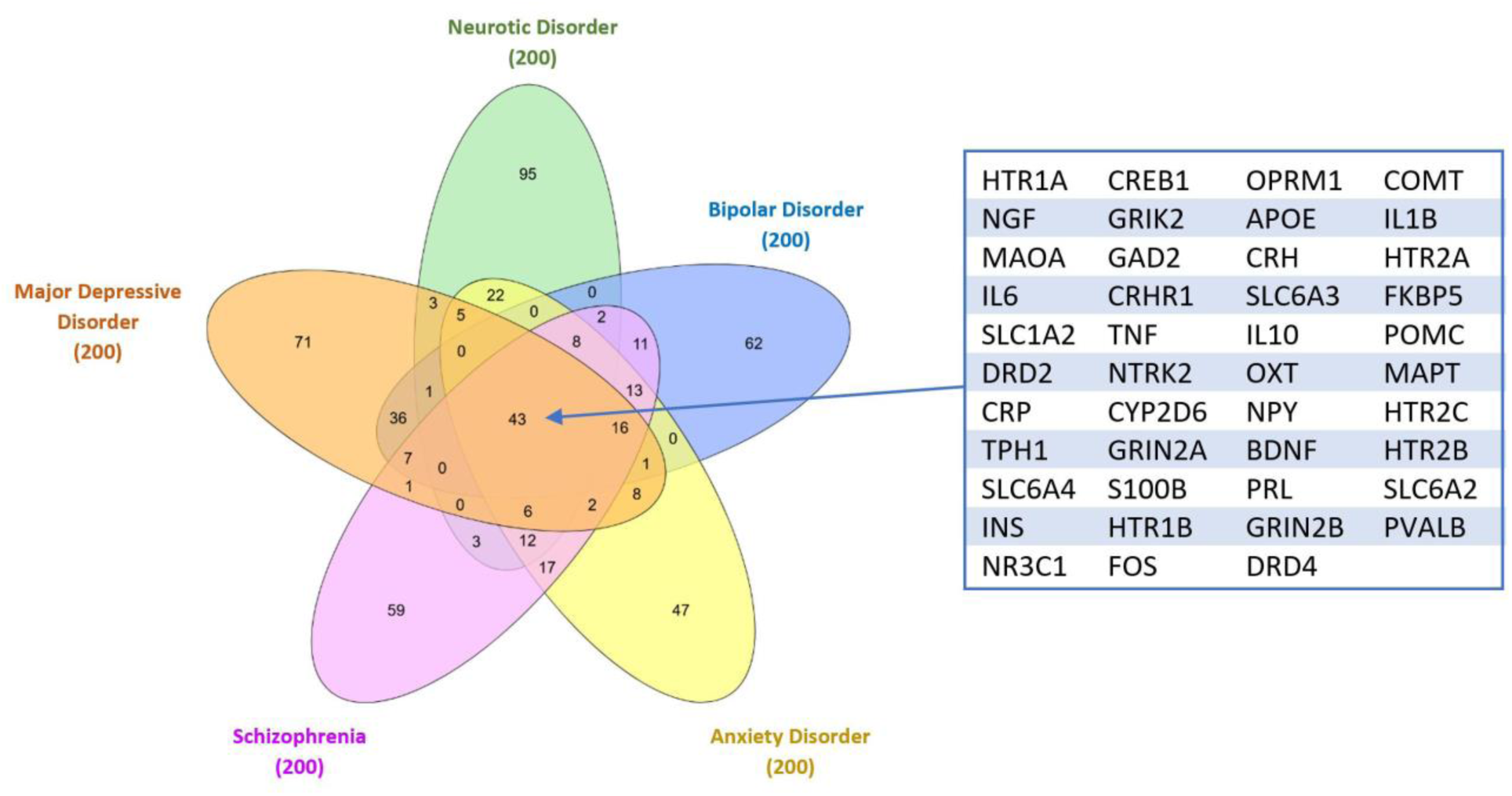
Comparison of the top 200 disease-associated proteins among the five NPDs (Schizophrenia, MDD, ND, AD and BD), indicating the presence of 43 common disease proteins.

**Table 2:** Major biological processes in which the common disease-associated proteins of NPDs participate.

| Biological Process | NPDs common disease-associated proteins |
| --- | --- |
| Serotonergic neurotransmission | 5-hydroxytryptamine receptor 1A (HTR1A), 5-hydroxytryptamine receptor 2A (HTR2A), Tryptophan 5-hydroxylase 1 (TPH1), Sodium-dependent serotonin transporter (SLC6A4), 5-hydroxytryptamine receptor 1B (HTR1B), 5-hydroxytryptamine receptor 2C (HTR1C) |
| Dopaminergic neurotransmission | Monoamine oxidase A(MAOA), D2 dopamine receptor (DRD2), Dopamine D4 receptor (DRD4) |
| Glutamatergic neurotransmission | Glutamate receptor ionotropic, kainate 2 (GRIK2), Glutamate receptor ionotropic NMDA 2A (GRIN2A), Glutamate receptor ionotropic, NMDA 2B (GRIN2B), Glutamate decarboxylase 2 (GAD2) |
| Inflammation | Corticotropin releasing hormone (CRH), Interleukin 10 (IL10), Tumor Necrosis Factor (TNF), Corticotropin releasing hormone receptor 1 (CRHR1), Interleukin-1 beta (IL1B), Glucocorticoid receptor (NR3C1) and Interleukin-6 (IL6) |
| Neurodegeneration | Apolipoprotein E (APOE), Microtubule-associated protein (MAPT) |
| Diabetes | Insulin (INS) |

### Analysis Results of the DM2 ∩ NPDs KEGG Pathway-to-Pathway Network

We reconstructed and analyzed the DM2 ∩ NPDs KEGG pathway-pathway network (see **Figure 3**), which consists of 328 functional relationships (edges) between the 87 common pathways (nodes) identified through our analysis to be shared between DM2 and all NPDs.

**Figure 3:**
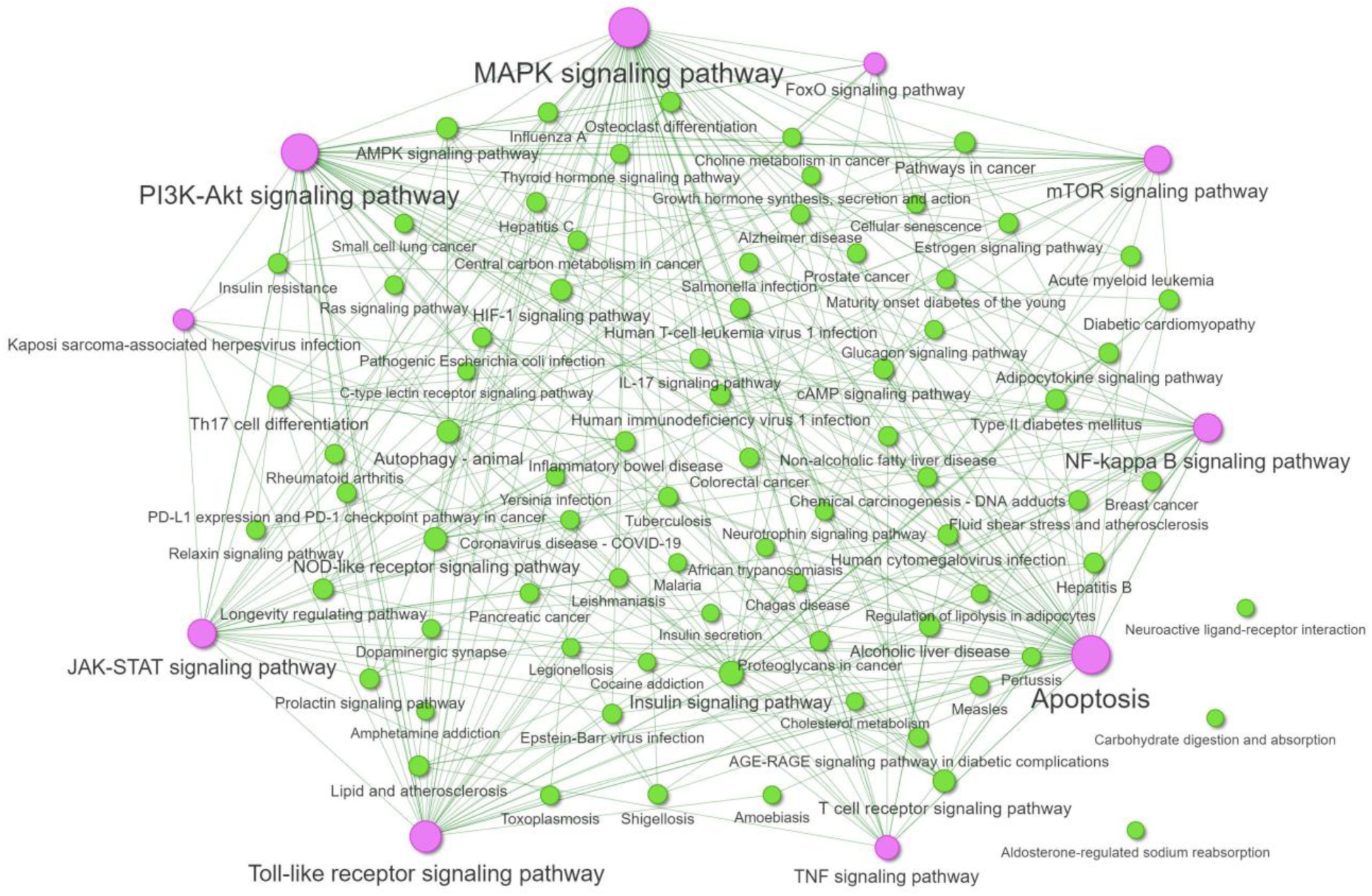
Visualization of the DM2 ∩ NPDs KEGG pathway-pathway network, indicating the functional relationships between the 87 common pathways between DM2 and NPDs. The top 10 hubs are highlighted in purple color. The size of the nodes is proportional to their degree.

#### (a) Composition of the DM2 ∩ NPDs KEGG Pathway-Pathway Network

The composition analysis of the DM2 ∩ NPDs KEGG pathway-pathway network indicated that the 87 common pathways belong to 21 subclasses (**Table 3**), according to the KEGG database classification system. The results revealed that 23 pathways belong to the subclass of infectious diseases (viral, bacterial and parasitic), while 10 pathways belong to the endocrine system subclass. In addition, 11 pathways were classified under the signal transduction subclass, and 12 pathways were associated with various cancer subclasses.

**Table 3:** Top 10 subclasses found in the DM2 ∩ NPDs KEGG pathway-pathway network and the number of pathways in each subclass.

| Rank | Subclass | Frequency (pathways) |
| --- | --- | --- |
| 1. | Signal transduction | 11 |
| 2. | Endocrine system | 10 |
| 3. | Infectious disease: viral | 10 |
| 4. | Infectious disease: bacterial | 7 |
| 5. | Cancer: overview | 6 |
| 6. | Cancer: specific types | 6 |
| 7. | Endocrine and metabolic disease | 6 |
| 8. | Immune system | 6 |
| 9. | Infectious disease: parasitic | 6 |
| 10. | Cardiovascular disease | 3 |

#### (b) Comorbidity (DM2 ∩ NPDs) Disease Communicator Nodes

Topological analysis of the network led to the identification of the top 10 hubs nodes (**Figure 3**, **Table 4**), which are high-centrality nodes that communicated with several of the other common disease pathways in the network. Therefore, these pathways, acting as potential comorbidity disease communicator nodes, are hypothesized to play a crucial role in facilitating the comorbidity between DM2 and NPDs. Their centrality in the network suggests they are essential due to their high connectivity and potential systemic influence.

**Table 4:** Top 10 high centrality nodes that facilitate the emergence of DM2 and NPD comorbidity and their classification.

| Rank | KEGG pathway name | Classification |
| --- | --- | --- |
| 1. | Apoptosis (hsa04210) | Cell growth and death |
| 2. | PI3K-Akt signaling pathway (hsa04151) | Signal transduction |
| 3. | MAPK signaling pathway (hsa04010) | Signal transduction |
| 4. | Toll-like receptor signaling pathway | Immune system |
| 5. | NF-kappa B signaling pathway (hsa04064) | Signal transduction |
| 6. | JAK-STAT signaling pathway (hsa04630) | Signal transduction |
| 7. | TNF signaling pathway (hsa04668) | Signal transduction |
| 8. | mTOR signaling pathway (hsa04150) | Signal transduction |
| 9. | Kaposi sarcoma-associated herpesvirus infection (hsa05167) | Infectious disease: viral |
| 10. | FoxO signaling pathway (hsa04068) | Signal transduction |

The rationale behind identifying ‘communicator nodes’ within our network stems from the need to prioritize and understand the most influential pathways among the numerous shared ones. While all 87 pathways identified contain genes associated with both DM2 and NPDs, network analysis allows us to pinpoint pathways that are central and highly connected, suggesting they might have a more significant systemic impact. This approach helps in prioritizing pathways for further study and potential therapeutic targeting. Moreover, network analysis provides a structured way to hypothesize about the functional relationships and interactions between pathways, which might not be apparent from gene overlap alone. The identification of high centrality pathways as potential ‘communicator nodes’ offers a focused direction for subsequent experimental validation and therapeutic exploration.

According to KEGG database classification system, seven of the communicator nodes belong to the subclass of ‘Signal transduction’ (**Table 4**). In contrast, Apoptosis belongs to the subclass of ‘Cell growth and death’, the Kaposi sarcoma-associated herpesvirus (KSHV) infection pathway belongs to the subclass of ‘Infectious disease: viral’, and the Toll-like receptor signaling pathway belongs to the ‘immune system’ subclass.

#### (c) Tissue Specificity Analysis Results of DM2-NPDs Interactions

Comorbid conditions often involve multiple tissues and organs. Therefore, considering the interplay of molecular changes in different tissues and their contributions to comorbidity is essential. To address this, we conducted tissue-specific enrichment analysis of the disease-associated genes from DM2 and NPDs that were found to participate in the 87 common disease pathways. This analysis aimed to identify potential overlapping tissues where the dysregulation of these pathways could occur between these two conditions. The tissue-specific enrichment analysis revealed 14 statistically significant tissues related to the disease-associated genes in DM2 (**Figure 4A**) and 17 statistically significant tissues in NPDs (**Figure 4B**). This approach highlighted 11 overlapping tissues (spleen, pancreas, cervix uteri, fallopian tube, colon, pituitary, small intestine, esophagus, stomach, muscle, brain) between DM2 and NPDs where the dysregulation of the 87 common disease pathways could occur.

**Figure 4:**
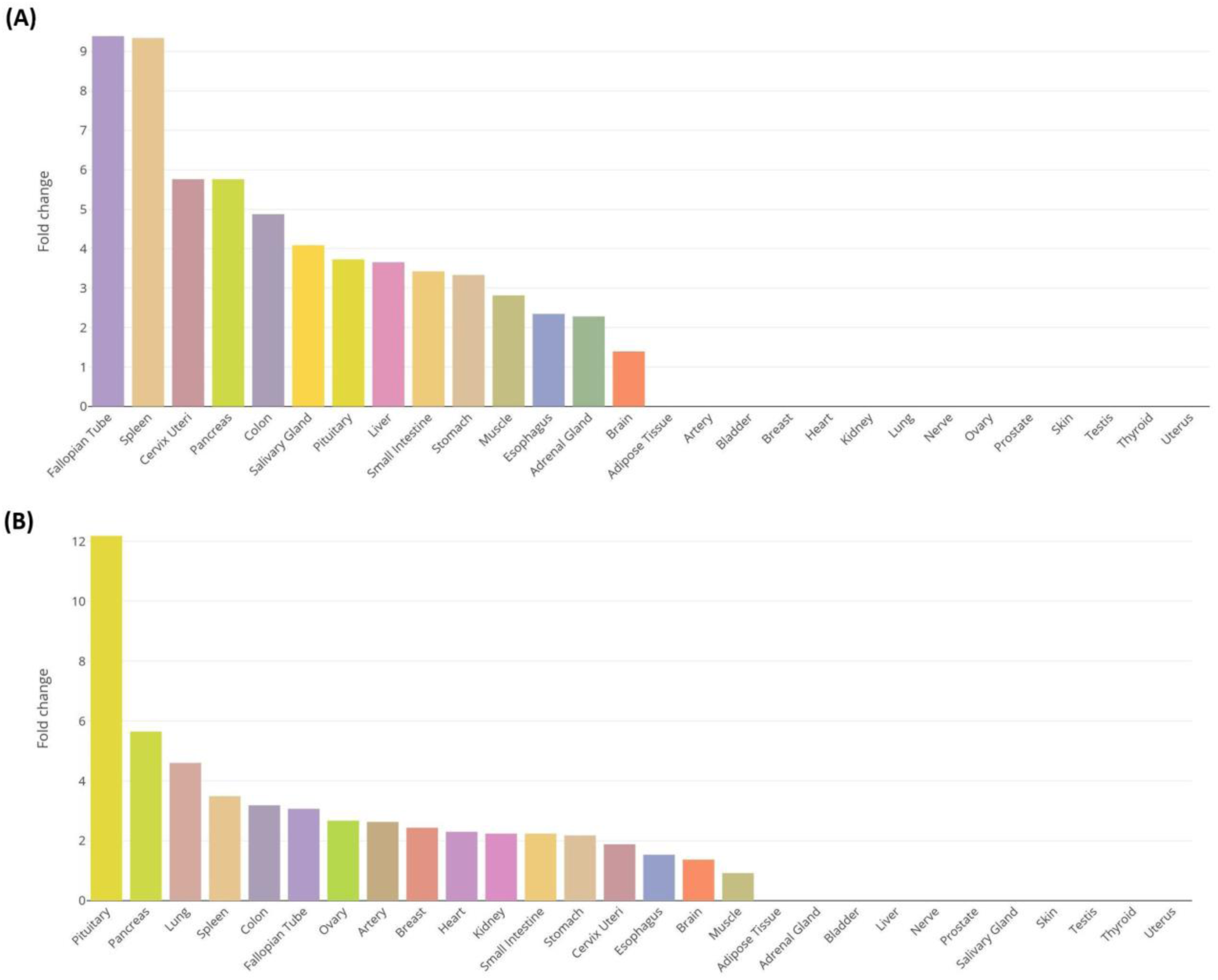
Tissue enrichment analysis results of disease-associated genes from **(A)** DM2 and **(B)** NPDs, which participate in the 87 common disease pathways.

### Isolating the Shortest Path to DM2 and NPDs Comorbidity

#### (a) Adding Reference Points and Missing Pathways on the DM2 ∩ NPDs KEGG Pathway-Pathway Network

Our objective was to identify the most critical path that might facilitate the development of comorbid DM2 and NPDs. To achieve this, we utilized the DM2 ∩ NPDs KEGG pathway-pathway network, which comprised the 87 common disease pathways identified between DM2 and NPDs. We selected five pathways to act as reference points for the two conditions (see Methods) and highlighted them on the DM2 ∩ NPDs KEGG pathway-pathway network. We also employed the shortest path approach (see Methods) to add any missing reference points and ensure connectivity within the network. In addition, to create a fully connected network, we identified nodes with degree values of 0 and 1, and employed the shortest path approach to add additional missing pathways to the network (see Methods) to generate a fully connected expanded DM2 ∩ NPDs KEGG pathway-pathway network (**Figure 5**), consisting of 97 nodes and 391 edge interactions. The remaining unconnected pathway lacked any direct functional interactions based on the data collected from the KEGG database (**Figure 5**). Additionally, our approach introduced seven additional pathways (highlighted in red) to establish a fully connected network. Our model highlights the significance of the added Calcium signaling pathway (hsa04020) in facilitating the development of comorbid DM2 and NPDs, as it exhibits functional relationships with several common disease pathways (DM2 ∩ NPDs) identified between DM2 and NPDs (**Figure 5**).

**Figure 5:**
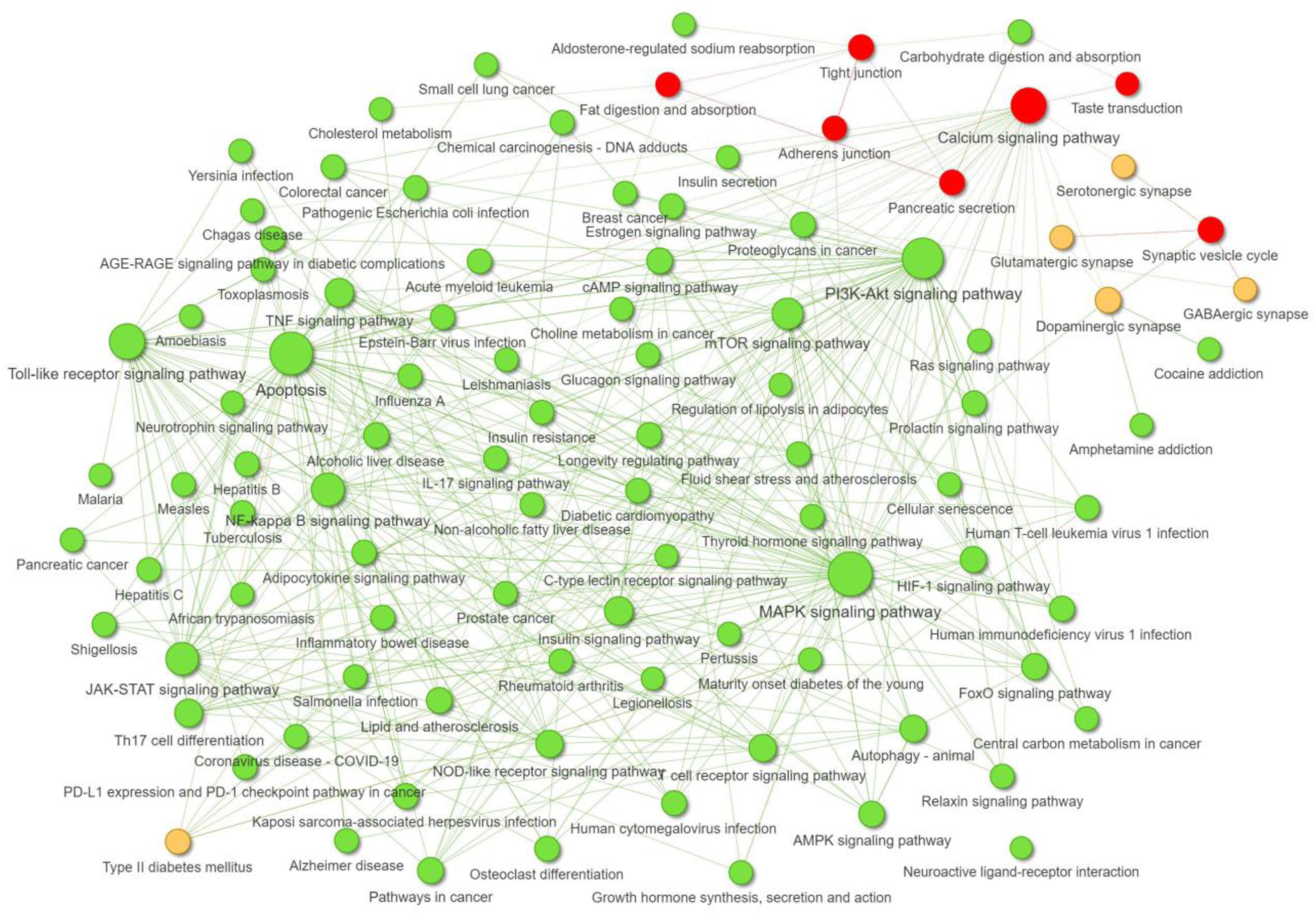
Visualization of the expanded DM2 ∩ NPDs KEGG pathway-pathway network, where the reference points are highlighted in yellow, the additional pathways added are shown in red, and the remaining common disease pathways between DM2 and NPDs are depicted in green.

#### (b) Highlighting the Shortest Path to DM2 and NPDs Comorbidity

To isolate the shortest path that contributes to the development of comorbid DM2 and NPDs, we initially calculated and isolated all the shortest paths with the smallest length between the DM2 reference point (Type II diabetes mellitus) and each of the four pathways representing NPDs (Dopaminergic synapse, Glutamatergic synapse, Serotonergic synapse and GABAergic synapse). Subsequently, we extracted the relevant nodes and edges from these shortest paths and highlighted them on the expanded DM2 ∩ NPDs KEGG pathways network (**Figure 5**), which allowed to isolate the shortest path that might lead to the emergence of DM2 and NPDs comorbidity.

The shortest path leading to NPDs and DM2 comorbidity encompasses three high centrality pathways: Calcium signaling pathway, MAPK signaling pathway and Apoptosis (**Figure 6**). These pathways not only exhibit functional interactions with one another but also have the potential to exert systemic comorbid pathogenic effects within the expanded DM2 ∩ NPDs KEGG pathway-pathway network (**Figure 5**). Therefore, dysregulation of these pathways can facilitate the development of comorbid DM2 and NPDs. Notably, the MAPK signaling and Apoptosis pathways are high centrality nodes in the initial DM2 ∩ NPDs KEGG pathway-pathway network. Moreover, the Calcium signaling pathway was an additional missing node added on the network to establish connections between the unconnected NPDs reference points and the rest of the network. It also interacts with several of the common disease pathways. In addition, the shortest path to comorbidity includes the Estrogen signaling pathway, which exhibits functional interactions with the GABAergic synapse (NPDs reference point), the Calcium signaling pathway (added missing pathway), and the MAPK signaling pathway (comorbidity disease communicator nodes).

**Figure 6:**
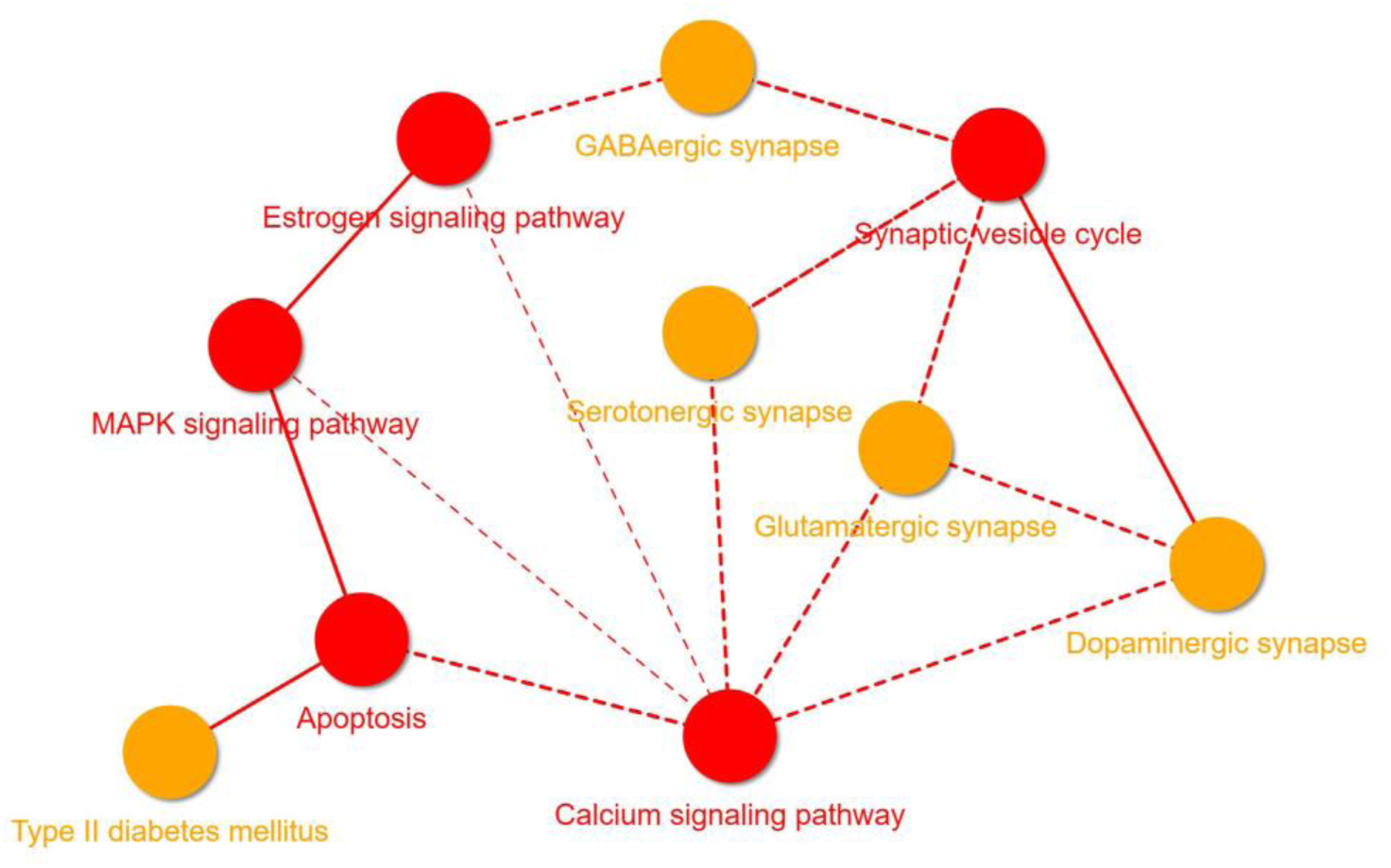
Shortest path to DM2 and NPDs comorbidity. Illustration of the subnetwork representing the shortest path to DM2 and NPDs comorbidity. It highlights the five reference points representing DM2 and NPDs in yellow color, as well as the minimum number of pathways (shown in red) required to connect the Type II diabetes mellitus pathway (representing DM2) with each of the four reference points: Dopaminergic synapse, Glutamatergic synapse, Serotonergic synapse and GABAergic synapse (representing NPDs). Dotted edges represent additional interactions that were required to be added to create the fully connected expanded DM2 ∩ NPDs KEGG pathway-pathway network. On the other hand, undotted edges depict the interactions present in the initial DM2 ∩ NPDs KEGG pathway-pathway network, which included only the 87 common disease pathways found to be shared between DM2 and NPDs.

### Validation of the importance of the central and “shortest path to comorbidity” pathways in DM2 and NPDs comorbidity using PFC Data

As described in the Methods section, we validated the importance of the central and “shortest path to comorbidity” pathways using microarray gene expression data from the PFC obtained from the GEO database.

Pathway enrichment analysis of the DEGs from each dataset revealed 8, 21, 13, and 28 statistically significant KEGG pathways associated with depression, schizophrenia, bipolar disorder (BD), and DM2, respectively (**Supplementary File 2-3**). The pathway enrichment analysis yielded several significant findings. First, the PI3K-Akt signaling pathway, identified as one of the top 10 high centrality nodes (hubs) in our computational model contributing to DM2 and NPDs comorbidity, was statistically significant in all three NPDs datasets (depression, schizophrenia, BD). Additionally, it showed statistical significance in the DEGs from the PFC of DM2 rats. Second, the Kaposi sarcoma-associated herpesvirus infection pathway, which was recognized by our analysis as one of the top 10 high centrality disease communicator nodes, displayed statistical significance in both DM2 and schizophrenia. Finally, the MAPK signaling pathway, highlighted in our analysis as participating in the shortest path to DM2 and NPDs comorbidity and as one of the top 10 high centrality nodes, demonstrated statistical significance in both NPDs (schizophrenia, BD) and DM2.

Previous studies have successfully used transcriptomics to infer shared and distinct molecular mechanisms underlying comorbid diseases, providing valuable insights into the pathophysiological links and potential therapeutic targets for complex comorbidities (61). The findings from the PFC transcriptomic datasets (GSE34451, GSE12654) indicate a strong convergence between the pathways identified through our network-based computational framework and those deemed statistically significant in the DEGs from the PFC. This convergence validates the reliability and robustness of our computational model by demonstrating that it can independently capture relevant pathways. This alignment reinforces the significance of these pathways in the comorbidity of DM2 and NPDs and confirms our model as a robust alternative for uncovering critical pathological mechanisms. Despite not being tissue-specific, our model captures critical interactions and offers a comprehensive understanding that is well-supported by PFC-specific data, highlighting its effectiveness and reliability in studying comorbid diseases. This validation process provides strong evidence supporting our computational findings and underscores the robustness of our pathway network-based approach in identifying key pathological mechanisms underlying the comorbidity of DM2 and NPDs.

## Discussion

The presence of comorbid diseases is associated with complicated treatment requirements, poorer prognosis, and increased healthcare costs (1, 2). Therefore, understanding the exact pathophysiological mechanisms that facilitate the emergence of comorbid diseases is essential for identifying suitable pharmacological treatments. Currently, there is a lack of transcriptomic data from human patients, as well animal models, with comorbid DM2 and NPDs available in well-known public transcriptomic repositories, such as ArrayExpress (62) and GEO (56). In this work, we developed and applied a network-based computational framework that offers an alternative and valid approach to overcome this limitation, serving as a robust alternative to traditional transcriptomic data analysis. By using our computational framework, which utilizes two complementary identification methods that operate synergistically, we were able to identify the top 10 high centrality (hub) disease pathways and the shortest path to comorbidity, whose dysregulation can facilitate the emergence of comorbid DM2 and NPDs. This methodological approach opens new avenues for understanding complex disease interactions and can potentially lead to the identification of novel therapeutic targets.

Our approach does not aim to capture all possible mechanisms that contribute to the emergence of this comorbidity but rather aims to highlight key pathways that lead to its emergence, mediated via the presence of co-shared disease pathways. The ability of our computational framework to identify key pathways associated with comorbidities, in this case DM2 and NPDs, was validated using transcriptomic data from the PFC.

We first reconstructed five DM2-NPDs comorbidity PPI networks and extracted their commonalities subnetworks. We then performed enrichment analysis on the human proteins contained in each subnetwork. This allowed to identify 87 common disease pathways shared between DM2 and all five NPDs (ND, MDD, BD, AD and Schizophrenia), which could represent possible crossroad pathological mechanisms that facilitate the emergence of comorbid DM2 and NPDs. Furthermore, we reconstructed the DM2 ∩ NPDs KEGG pathway-pathway network using the identified 87 common disease pathways and their 328 functional relationships.

Topological analysis of the DM2 ∩ NPDs KEGG pathway-pathway network allowed us to identify the top 10 high centrality (hubs) pathways, which can act as comorbidity disease communicator nodes and exert systemic comorbid pathogenic effects, hence have a key role in facilitating the emergence of this comorbidity. Nodes that act as hubs in a network have a high degree of connectivity, meaning they are connected to a large number of other nodes in the network, this centrality renders them crucial for the network’s overall structure and functioning. Consequently, perturbation introduced to hub nodes can have significant implications for the network’s stability, functioning, and overall behavior.

Our network analysis identified the top 10 high centrality pathways that could possibly lead to DM2 and NPDs comorbidity, suggesting their essential role due to their high connectivity and potential systemic influence. These pathways, including PI3K-Akt signaling, mTOR signaling, and Toll-like receptor signaling pathway, which have been previously implicated in the pathophysiology of both DM2 and NPDs. For instance, the PI3K-Akt signaling pathway is crucial for insulin signaling and glucose homeostasis and is known to be dysregulated in DM2 (63). Dysregulation of this pathway in the brain has been associated with synaptic dysfunction and neuroinflammation (64), which are critical factors in the development of NPDs such as schizophrenia and depression. The mTOR signaling pathway, another high centrality node, plays a vital role in cell growth, proliferation, and survival. Alterations in mTOR signaling have been associated with both metabolic disorders like DM2 (63) and NPD conditions such as MDD (65, 66), BD (67) and Schizophrenia (68). Evidence from preclinical models suggests that impaired mTOR signaling contributes to insulin resistance and β-cell dysfunction in DM2 and affects synaptic plasticity and neurogenesis in NPDs (63, 65–68). Chronic inflammation is a common feature in DM2 and has been increasingly recognized in the pathology of NPDs, including the possible role of Toll-like receptor signaling in contributing to this inflammation (69–71). The identification of immune-related pathways suggests that inflammatory processes may be a shared mechanism contributing to the comorbidity.

We have also analyzed the composition of the 87 common disease pathways, which showed that the majority belong to the subclasses of Signal transduction, Endocrine system, Cancer, and Infectious diseases. More specifically, 23 pathways belong to the infectious disease subclasses, including viral, bacterial, and parasitic infections. Several pathogenic organisms have been considered as environmental risk factors for the development of various diseases, including DM2 (72–77) and NPDs (78, 79). Based on the ‘associated risk factors’ etiological model, comorbidity can arise when risk factors, such as viral infections, for Disease “A” correlate with the risk factors of Disease “B” (1, 80, 81). For example, the COVID-19 infectious disease is associated with the development of both DM2 and several NPDs, including depression, schizophrenia, and anxiety disorders (31, 74, 75, 79, 82). Viruses have the ability to modulate and dysregulate disease-associated pathways via virus-host PPIs and lead to the emergence of diseases (31, 34, 35), thereby leading to the development and exacerbation of comorbid conditions. Understanding these mechanisms highlights the importance of infectious disease pathways in contributing to the comorbidity of DM2 and NPDs, emphasizing the need for integrated therapeutic strategies that address both metabolic and neuropsychiatric aspects.

In addition, we performed tissue-specific gene enrichment which led to the identification of eleven overlapping tissues (spleen, pancreas, cervix uteri, fallopian tube, colon, pituitary, small intestine, esophagus, stomach, muscle, brain) between DM2 and NPDs. These tissues could represent possible sites where the dysregulation of the 87 common pathways could occur. The pancreas and the brain are particularly notable, as their reciprocal interaction plays an important role in maintaining glucose homeostasis both in the brain and peripheral tissues (83). Pancreatic islets communicate with the brain, and *vice versa*; brain circuits regulate the endocrine functions of the pancreas. The brain is a key player in the regulation of energy metabolism and glucose homeostasis, as it integrates various peripheral metabolic inputs, including signals from the pancreas (84). Glucose metabolism is critical for brain functioning, and disruption of glucose metabolism is a primary pathophysiological characteristic of NPDs (85). Hence, the brain is particularly vulnerable to the metabolic effects of DM2. Moreover, evidence suggests the involvement of brain pathways in the regulation of pancreatic islet physiology; however the exact brain regions that communicate with the pancreas have not yet been fully defined (86, 87). Therefore, communication between common disease pathways from different tissues can also facilitate the development of comorbidities.

It is notable that the fallopian tube appeared as the most enriched tissue in our analysis of DM2 (Figure 4A). This unexpected result underscores the need for further investigation to understand its relevance. One possible explanation is the association between DM2 and polycystic ovarian syndrome (PCOS) in women. Women of reproductive age who have PCOS and are obese have an eight times greater chance of developing DM2 (88). This is because most PCOS women exhibit metabolic syndrome symptoms such as insulin resistance (89). Additionally, women with PCOS have a higher prevalence of NPDs, including anxiety, depression, and BD (90). This link suggests that molecular changes in the fallopian tube related to PCOS might also be relevant in the context of DM2 and NPDs comorbidity.

Finally, we devised the ‘minimum path to comorbidity’ algorithm that allowed us to identify the shortest path that might facilitate the development of comorbid DM2 and NPDs. Existing tools utilizing different methods allow for the recreation, visualization, and analysis of pathway-to-pathway networks, such as ComPath (91), ClueGO (43), PANEV (92) and PathExNET (93). While some of these tools allow to introduce complementary pathways to create a fully connected network (94), our approach allows for the introduction of specific pathways into the network that act as disease reference points, making them relevant to the diseases under investigation. It also allows for the selection of specific pathways to represent the disease, even in cases where no available disease pathways are present in the KEGG database. In this particular case, the NPDs pathways were not available in the KEGG database, so we selected four relevant pathways to represent the NPDs pathway.

Most importantly, our novel approach allows for the highlighting of all the shortest paths on a KEGG pathway-to-pathway network between selected pathways that act as disease reference points. This feature enables us to highlight the minimum path that might facilitate the development of comorbid diseases. The ‘minimum path to comorbidity’ algorithm is designed to identify the most direct and functionally relevant connections between disease pathways that contribute to the comorbidity of DM2 and NPDs. While it leverages the shortest path principle, it integrates biological relevance by focusing on pathways with established roles in both conditions. This targeted approach ensures that the identified paths are not only the shortest but also biologically significant. Therefore, the ‘minimum path to comorbidity’ algorithm is not merely a shortest path algorithm but a biologically informed method that integrates network centrality, functional relevance, and enrichment validation. This approach provides meaningful insights into the pathophysiological mechanisms underlying the comorbidity of DM2 and NPDs and identifies potential therapeutic targets.

The ‘minimum path to comorbidity’ algorithm allowed us to highlight the Calcium signaling pathway, MAPK signaling pathway, Apoptosis pathway, and the Estrogen signaling pathway as pathways within the shortest path that leads to DM2 and NPDs comorbidity. The identification of these pathways suggests their critical role in the interplay between DM2 and NPDs. The fact that the MAPK signaling pathway and Apoptosis pathway are also ranked among the top 10 high centrality nodes within the DM2 ∩ NPDs KEGG pathway-pathway network further supports their central role in the emergence of comorbid DM2 and NPDs. These findings suggest that pharmacological interventions targeting these pathways may offer a promising approach to simultaneously address both diseases, due to their close proximity to the reference points of DM2 and NPDs and their functional interactions. Despite the existence of other possible paths, our hypothesis is that targeting these pathways would have a higher drug impact effect than targeting more distant disease pathways or longer paths involving additional pathways. This hypothesis is supported by network-based approaches that have modeled the effects of drugs and have shown that proximity between drug targets and the disease pathways provides new insights into the therapeutic effects of pharmacotherapies (95, 96).

Calcium signaling plays an important role in the regulation of insulin secretion from pancreatic islet β-cells (97) and its dysregulation results in deficient insulin secretion, increasing the risk for the development of DM2 (98). In turn, deficient pancreatic insulin release triggers calcium dysregulation in neuronal cells, leading to the impairment of synaptic plasticity in the brain (99) and neuronal cell death (100, 101) which facilitates the development of brain diseases, including NPDs, as well as neurodegenerative diseases (NDs) like Alzheimer’s disease and Parkinson’s Disease. Thus, calcium signaling is essential for both insulin secretion and neuronal function, and its dysregulation can lead to metabolic and cognitive impairments. Additionally, calcium signaling can promote neuronal apoptosis via the activation of the MAPK signaling pathway (102, 103). The MAPK signaling pathway also activates various transcription factors involved in learning and memory, and abnormal activity of the MAPK signaling pathway contributes to the development of NPDs (104, 105). The MAPK signaling pathway is a key signal transduction pathway that plays an important role in the regulation of various physiological functions, including insulin signaling, and its improper activation also plays a role in diabetic complications (106).

Moreover, estrogen signaling has neuroprotective effects and regulates glucose metabolism, with low estrogen levels being linked to increased risk of both DM2 and NPDs. Estrogen, which is a sex hormone, exerts neuroprotective effects in the brain, through activation of the MAPK signaling pathway (107–109). Thus, low estrogen levels in the brain result in reduced activation of the MAPK signaling pathway, diminishing the neuroprotective actions of estrogen, leading to brain diseases like NPDs and NDs (107). In addition, estrogen is an important regulator of glucose homeostasis, and low levels of estrogen in postmenopausal women are associated with an increased risk of developing DM2, insulin resistance, and decreased calcium secretion, which can be reversed with estradiol treatment (110, 111). In contrast to women, increased estradiol levels in men are associated with increased risk of developing DM2 (112). Due to the important functional relationships of the identified comorbidity pathways, they could represent potential pharmacological targets for the treatment of comorbid DM2 and NPDs. Additionally, estradiol’s impact on serotonin, glutamate, and dopamine systems (113) underscores the potential of targeting hormonal pathways to manage both DM2 and NPDs.

The highlighted pathways identified through our methodology represent highly promising pharmacological targets for the treatment and prevention of comorbid DM2 and NPDs. Moreover, our approach is grounded in sound biological knowledge, drawing on a wealth of existing research on DM2 and NPDs, and was validated utilizing publicly available transcriptomic experimental data. Overall, our study represents a major advance in the field of comorbid disease research and has significant potential to impact clinical practice by enabling the development of more effective treatments for this complex and challenging comorbid condition. Our methodology can serve as a paradigm for identifying key pathological mechanisms underlying the emergence of other comorbid diseases, making it a valuable resource for researchers and clinicians alike.

## Data Availability

The data used in this article are derived from publicly available sources. The microarray GSE34451 and GSE12654 used for this study can be found in the Gene Expression Omnibus database (https://www.ncbi.nlm.nih.gov/geo/).

Additional data used in this study were:

KEGG database (https://www.genome.jp/kegg/pathway.html);

*STRING disease* app (https://apps.cytoscape.org/apps/stringapp);

DISEASES database (https://diseases.jensenlab.org/Search)

## Code Availability

All of the programs we use in this study for analysis are free and open-access software.

Information on the version of the open-source software and its official website:

1. Cytoscape (v3.8.2) https://cytoscape.org/
2. stringAPP (v2.0.1) https://apps.cytoscape.org/apps/stringapp
3. ClueGO (v2.5.10) https://apps.cytoscape.org/apps/cluego
4. KEGGREST (v1.42.0) https://bioconductor.org/packages/release/bioc/html/KEGGREST.html
5. TissueEnrich web application(v1.22.0) https://tissueenrich.gdcb.iastate.edu/
6. Igraph R package (v1.5.1) https://igraph.org/

The software referenced in this study is open-access and can be obtained from their respective repositories. All other codes that support other findings of this study are available from the corresponding authors upon request.

## Authors contribution

A.O. and P.Z have contributed to the conceptualization, review and editing of the manuscript. A.O. developed the methodology, collected and analyzed the data, and wrote the original draft of the manuscript. All authors have read and agreed to the published version of the manuscript.

## Supplementary data

Supplementary File 1_ Reliability of the graph expansion approach

Supplementary File 2_ DEGs

Supplementary File 3_ Enriched terms

