## Supplementary File 1 for "Network-Based Analysis Identifies Targetable Pathways in Comorbid Type II Diabetes and Neuropsychiatric Disorders"

**Supplementary File 1: Reliability of the graph expansion approach**

One of the aims of our methodology is to identify complementary missing pathways that allow to interconnect pathways acting as disease reference points in a Disease ∩ Disease KEGG pathway-pathway comorbidity network with the rest of the pathways in the network. In this case, the DM2 and NPDs reference points were missing from the DM2 ∩ NPDs KEGG pathway-pathway network, which includes the 87 common disease pathways identified between DM2 and NPDs through enrichment analysis. These reference points needed to be added to the network using the shortest path method.

Additionally, our methodology aims to interconnect all pathways in the DM2 ∩ NPDs KEGG pathway-pathway network, including those with degree values of 0 or 1, with the other comorbidity pathways to establish connections between all the identified common disease pathways. To achieve this, our method introduces additional missing nodes using the shortest path method. These complementary missing pathways may not have been identified through enrichment analysis. Therefore, the complementary pathways could represent disease pathways associated with the emergence of these diseases, even if they were not identified as statistically significant in pathway enrichment analysis. This limitation is inherent to pathway enrichment analysis, as the standard approach is to select only statistically significant pathways. However, pathways that are not revealed as statistically significant could still be important, and our graph expansion method aims to overcome this limitation. Moreover, the complementary pathways may not have been associated with the development of these diseases in the current available literature or databases. Hence, validating our findings using currently available evidence may underestimate the importance of these new findings. Therefore, this approach also aims to identify new disease pathways involved in the emergence of this comorbidity.

To test the reliability of our graph expansion approach in introducing relevant disease pathways in the DM2 ∩ NPDs KEGG pathway-pathway network, we first compared the genes associated with each of the seven complementary missing pathways to the genes contained in the five pathways that act as reference points for DM2 and NPDs. In addition, we selected seven random pathways from the remaining 256 KEGG pathways that are not included in the expanded connected DM2 ∩ NPDs KEGG pathway-pathway network. We then determined the number of randomly selected pathways that have common genes with the reference point pathways of DM2 and NPD. We repeated this process of selecting seven random pathways and testing their gene commonalities with the reference points 30 times. In **Supplementary Figure 1**, the red line indicates that five out of the seven complementary missing pathways (as denoted by the red line), identified through our graph expansion algorithm, contain common genes with all five reference points. The green line represents the number of randomly selected pathways out of the seven that contain genes common with all five reference points. **Supplementary Figure 1** demonstrates that our graph expansion approach has a higher probability of introducing relevant pathways rather than selecting random ones.

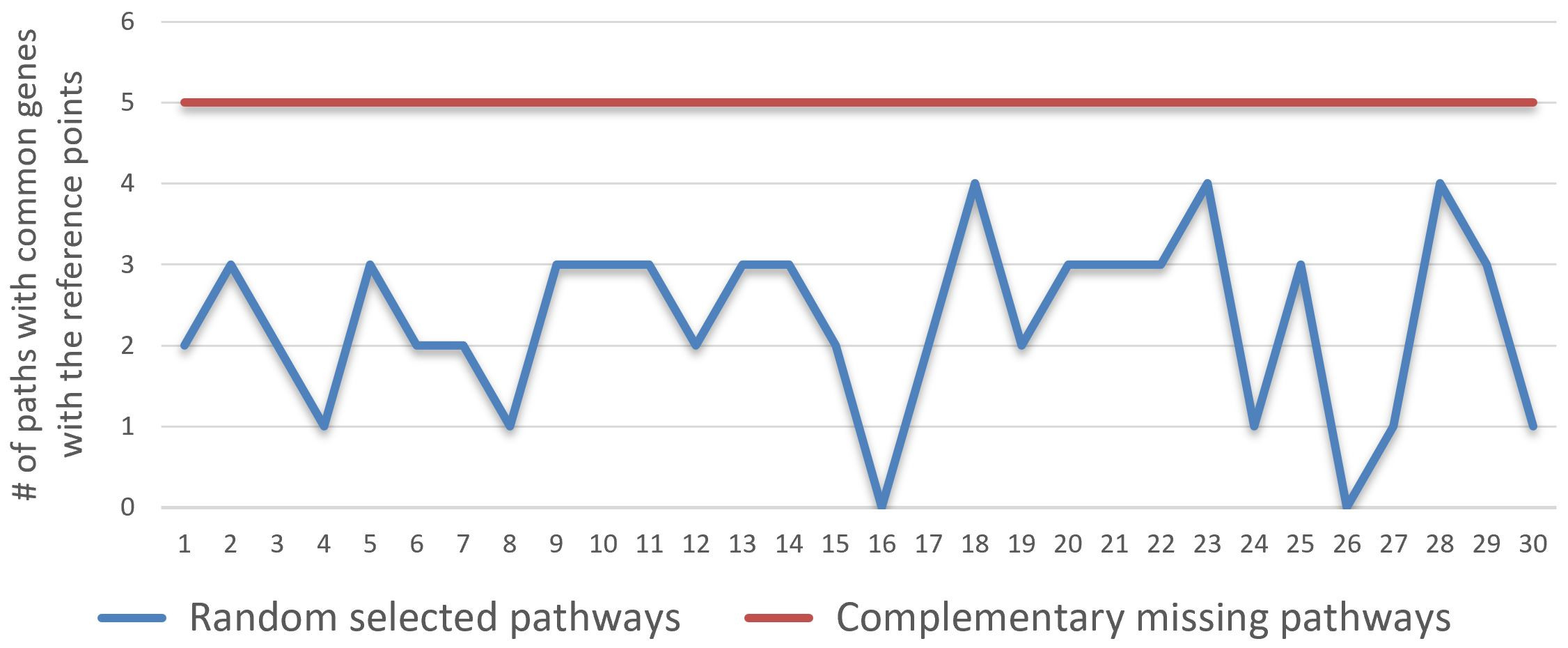

**Supplementary Figure 1:** The graph illustrates the ability of our graph expansion approach to introduce relevant pathways (red line) that have common genes with the reference points compared to randomly selected pathways (blue line).
